# One step 4x and 12x 3D-ExM: robust super-resolution microscopy in cell biology

**DOI:** 10.1101/2024.08.13.607782

**Authors:** Roshan X Norman, Yu-Chia Chen, Emma E Recchia, Jonathan Loi, Quincy Rosemarie, Sydney L Lesko, Smit Patel, Nathan Sherer, Motoki Takaku, Mark E Burkard, Aussie Suzuki

**Author notes:** Corresponding authors: Aussie Suzuki, Mark E Burkard. Equal contribution (Emma E Recchia, Jonathan Loi). Equal contribution (Roshan X Norman, Yu-Chia Chen).

## Abstract

Super-resolution microscopy has become an indispensable tool across diverse research fields, offering unprecedented insights into biological architectures with nanometer scale resolution. Compared to traditional nanometer-scale imaging methods such as electron microscopy, super-resolution microscopy offers several advantages, including the simultaneous labeling of multiple target biomolecules with high specificity and simpler sample preparation, making it accessible to most researchers. In this study, we introduce two optimized methods of super-resolution imaging: 4-fold and 12-fold 3D-isotropic and preserved Expansion Microscopy (4x and 12x 3D-ExM). 3D-ExM is a straightforward expansion microscopy method featuring a single-step process, providing robust and reproducible 3D isotropic expansion for both 2D and 3D cell culture models. With standard confocal microscopy, 12x 3D-ExM achieves a lateral resolution of under 30 nm, enabling the visualization of nanoscale structures, including chromosomes, kinetochores, nuclear pore complexes, and Epstein-Barr virus particles. These results demonstrate that 3D-ExM provides cost-effective and user-friendly super-resolution microscopy, making it highly suitable for a wide range of cell biology research, including studies on cellular and chromatin architectures.

## Introduction

In recent decades, fluorescence microscopy has emerged as an essential tool for pinpointing the locations, architectures, and dynamics of proteins and genes within cells. However, the resolution of conventional fluorescence microscopy is constrained to approximately 250 nm laterally and 500 nm axially due to its point spread function (PSF)^1,2^. This limitation means that many cellular macromolecular protein complexes and microbes, such as vertebrate kinetochores (∼250 nm)^3, 4^, microtubules (∼25 nm)^5–7^, nuclear pores (∼120 nm)^8, 9^, and viruses (∼100 nm)^10^, are smaller than the resolution limit of traditional fluorescence microscopy. To overcome this optical barrier, “super-resolution microscopy” has been developed, allowing researchers to study the structures, spatiotemporal dynamics, and functions of those nanoscale biomolecules with higher resolution^11–14^. Nonetheless, this advanced technique often necessitates specialized optical equipment, specific fluorescent dyes, or computational post-processing for image reconstruction, thereby limiting its widespread application^14^.

Expansion Microscopy (ExM) is a cutting-edge super-resolution microscopy technique that enhances resolution by physically expanding biological specimens, eliminating the need for expensive super-resolution microscopes^15^. In this method, cell cultures, organoids, and tissues are fixed and embedded into expandable hydrogel polymers. The gel-specimen composite is then expanded by absorbing water, resulting in enhanced resolution proportional to the expansion rate. The original and commonly used ExM achieves approximately 4-fold expansion^15^, theoretically reaching ∼60 nm lateral resolution with conventional light microscopy. Since many biological structures are smaller than this resolution limit, optimized ExM methods have been developed to achieve greater than 4-fold expansion. One approach involves sequential 4-fold expansion processes, which has been reported to achieve ∼20-fold expansion^16, 17^. However, the iterative expansion process has several limitations: complicated sample preparation, time-consuming, low reproducibility of expansion rate, and potential structural distortion. Another approach utilizes different gel chemistry, enabling ∼10-fold expansion, but requires specialized equipment to remove oxygen during gel polymerization^18^. The goal for the next generation of ExM is to achieve greater than 4-fold isotropic expansion with a simple single-step process.

Here, we introduce two robust ExM methods, 4x and 12x 3D-ExM, which ensure both 3D isotropic expansion and structural preservation of biospecimens. Researchers can choose either the 4-fold or 12-fold expansion protocol based on their desired resolution, and both involve single-step sample expansion without the need for specialized instruments or chambers. We validate the 3D isotropy of 3D-ExM by measuring the area and volume of nuclei, the largest organelle in mammalian cells, where achieving isotropic expansion has been challenging with previous ExM protocols^19, 20^. Additionally, we demonstrate that 3D-ExM resolves biological structures below the diffraction limit, including 1) the nuclear and cytoplasmic rings within a single nuclear pore complex, 2) individual viral particles of Epstein-Barr virus (EBV), 3) genomic RNA of human immunodeficiency virus (HIV), and 4) the human kinetochores during mitosis.

## Results

### Validation of 3D isotropic nuclear expansion by 3D-ExM

The workflow of 4x and 12x 3D-ExM is illustrated in **Fig. 1A**, with a detailed protocol in **Methods**. This protocol introduces several innovative steps, including the assembly of a home-made reusable imaging chamber that minimizes the drift of expanded hydrogels. 3D-ExM approach offers researchers two distinct hydrogel recipes, enabling either 4-fold (called 4x 3D-ExM) or 11∼12-fold (12x 3D-ExM) expansion in a single-step process. The 4x 3D-ExM hydrogel consists of acrylamide and N, N-methylenebisacrylamide (MBAA), as used in the original ExM method^15^. In contrast, the 12x 3D-ExM hydrogel is composed of N, N-dimethylacrylamide (DMAA) and sodium acrylate (SA), based on a protocol previously used for 10-fold gel expansion that required an oxygen-free environment^18^. We have developed robust DMAA-based polymerization technique that simplifies this process by using a piece of paraffin film to minimize air contact during the gel formation. This innovation enables easy and reproducible DMAA-based hydrogel polymerization without the need for special equipment. We confirmed that both MBAA- and DMAA-based hydrogels expanded isometrically by ∼4-fold and ∼12-fold, respectively, in both diameter and thickness compared to the pre-expanded gels (**Fig. 1B**). Note that full expansion requires 2-3 hours for the 4x hydrogel, whereas ∼20 hours are needed for the 12x hydrogel.

**Figure 1.**
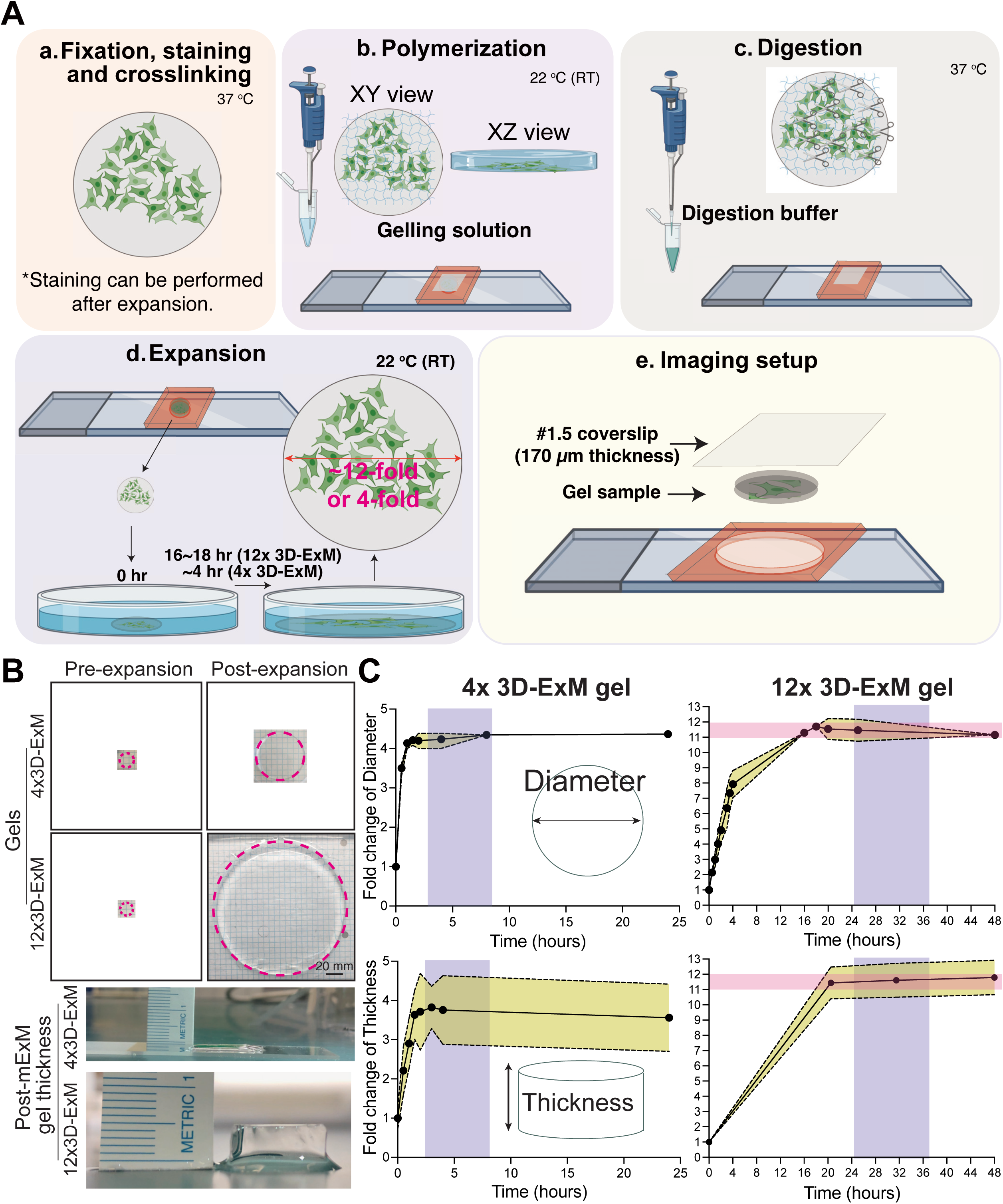

Next, we assessed the expansion isotropy of biological specimens embedded in the two types of hydrogels. Specifically, we focused on the nucleus of interphase cells, the largest and most structurally intricate organelle in mammalian cells. Previous ExM methods reported that the mammalian nucleus did not expand proportionally to gel expansion rates, resulting in anisotropy even along the xy-axis^19, 20^. We measured lateral expansion by averaging the lengths of the major and minor axes of the nucleus before and after expansion **(Fig. S1A)**. Axial expansion was determined by the volume of the nucleus through 3D rendered surface fitting (**Fig. S1B)**. We first evaluated nuclear expansion rates in 4x hydrogel and 12x hydrogel using the original digestion and crosslink protocols of 4xExM^15^ and 10xExM^18^ with cervical carcinoma HeLa cells and rat kangaroo PtK2 cells. Surprisingly, the nuclei of HeLa and PtK2 cells expanded only by 2.8∼2.9-fold in 4x ExM, and 4.5∼6.5-fold in 10xExM, despite both hydrogels robustly expanding by 4-fold and 12-fold, respectively (**Fig. 2A-B**). These findings suggest that the original ExM protocols do not achieve isotropic expansion for all cellular structures. Given that the chromatin fibers in the nucleus might restrict nuclear expansion, we treated the samples with micrococcal nuclease (MNase) before expansion. However, this did not improve the expansion of the nucleus, indicating that DNA linkage does not limit nuclear expansion (**Fig. S2A**). Next, we explored whether crosslinking between proteins and hydrogel polymers restricts nuclear expansion. We tested various concentrations of Acrylolyl-X (AcX), a protein crosslinking reagent commonly used in ExM^18, 21, 22^, and found that lower AcX concentrations yielded a higher nucleus expansion rate (**Fig. S2B**). This suggested that excessive chromatin crosslinking was the primary cause of limited expansion. However, the nuclear structure became distorted at low concentrations of AcX (< 30 µg/ml) (**Fig. S2B**). We then tested glutaraldehyde (GA) as an alternative crosslinker. We found that 0.05 - 2.1% GA alone allowed for 12-fold expansion, whereas GA combined with AcX failed to support 12-fold expansion (**Fig. S2C-D**). Notably, the nucleus could expand by ∼10-fold without any crosslinker, but the nucleus structure was significantly distorted (**Fig. S2D)**. These results indicate that AcX limits gel expansion and is not suitable for greater than 4x ExM. Although GA is known to cause autofluorescence, which typically requires quenching in the traditional staining protocols^23, 24^, we observed no detectable autofluorescence after expansion within the tested range of GA concentrations. We reasoned that the reduced autofluorescence in our ExM hydrogel-embedded biospecimens might be due to the formation of covalent bonds between GA and the gel matrix, which masked the aldehyde groups of GA.

**Figure 2.**
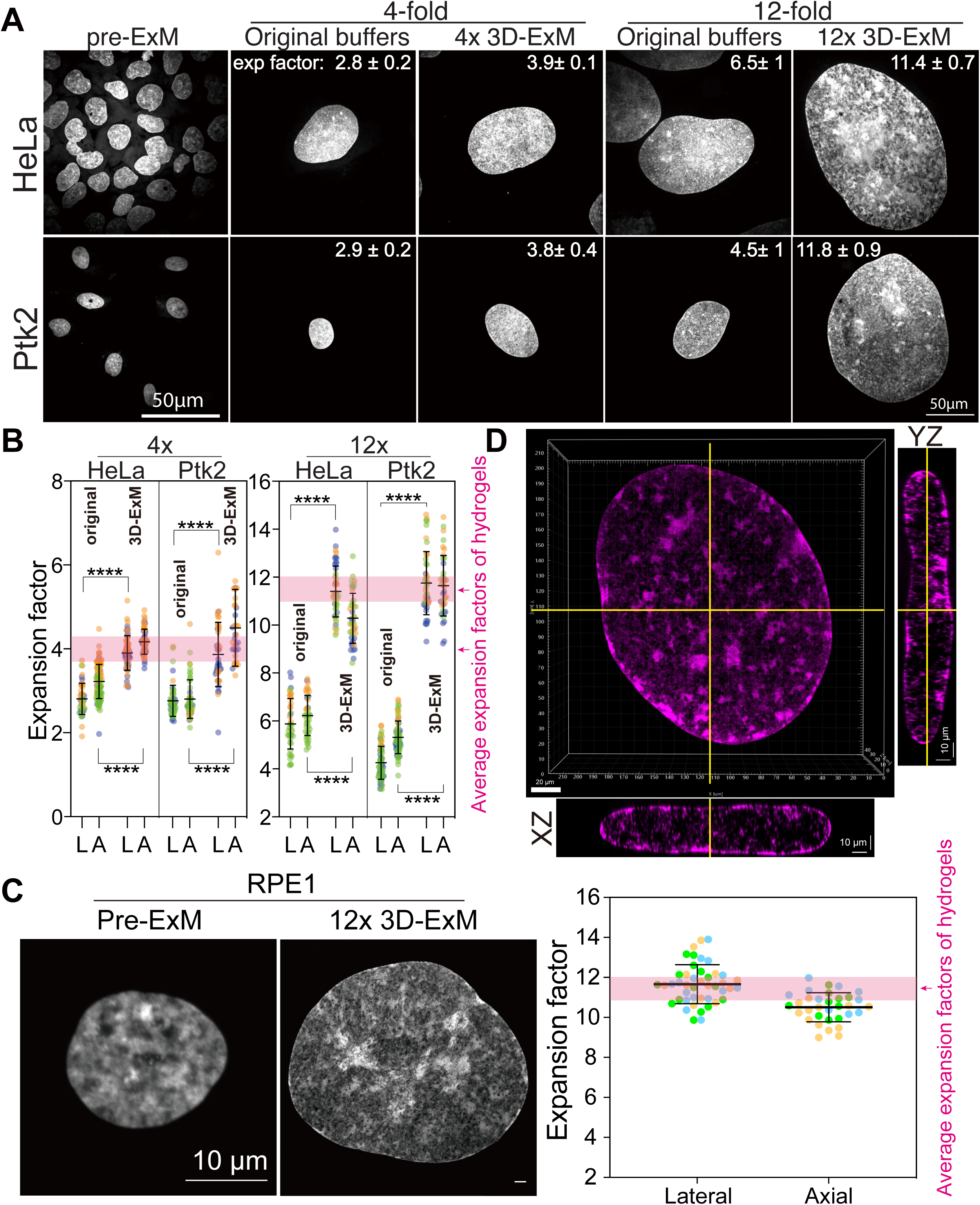

Using our modified protocol, which features optimized crosslinking and digestion steps, we achieved a successful expansion of interphase nucleus of HeLa, PtK2, and human retinal pigment epithelial (RPE1) cells by approximately 4-fold and 12-fold in three dimensions (**Fig. 2A-D**). Notably, the expanded interphase nuclei preserved typical heterochromatin domains, which are visible as regions with high DNA dye intensity (**Fig. 2D**). These results demonstrate that our 3D-ExM can robustly and isotropically expand nuclei within the 2D cell monolayer.

### Validation of Isotropic Expansion by Correlative Pre- and Post-3D-ExM

To further validate isotropic expansion in 4x and 12x 3D-ExM, we imaged the same cells before and after applying 4x or 12x 3D-ExM under identical imaging conditions (**Fig. 3A-D**). Initially, we imaged RPE1 cells with a 20x objective prior to expansion. After measuring the gel expansion rates, we imaged the same cells again using the same 20x objective. We calculated the expansion rates based on the lengths of long and short axes, as well as the area of the same cell before and after expansion. In both 4x and 12x 3D-ExM, the interphase nucleus expanded proportionally to the gel expansion rates, confirming equal and isotropic expansion between specimens and hydrogels. The nucleus size observed in 3D-ExM matched the digitally enlarged size based on the gel expansion factor, demonstrating its isotropic expansion and a significant improvement in resolution.

**Figure 3.**
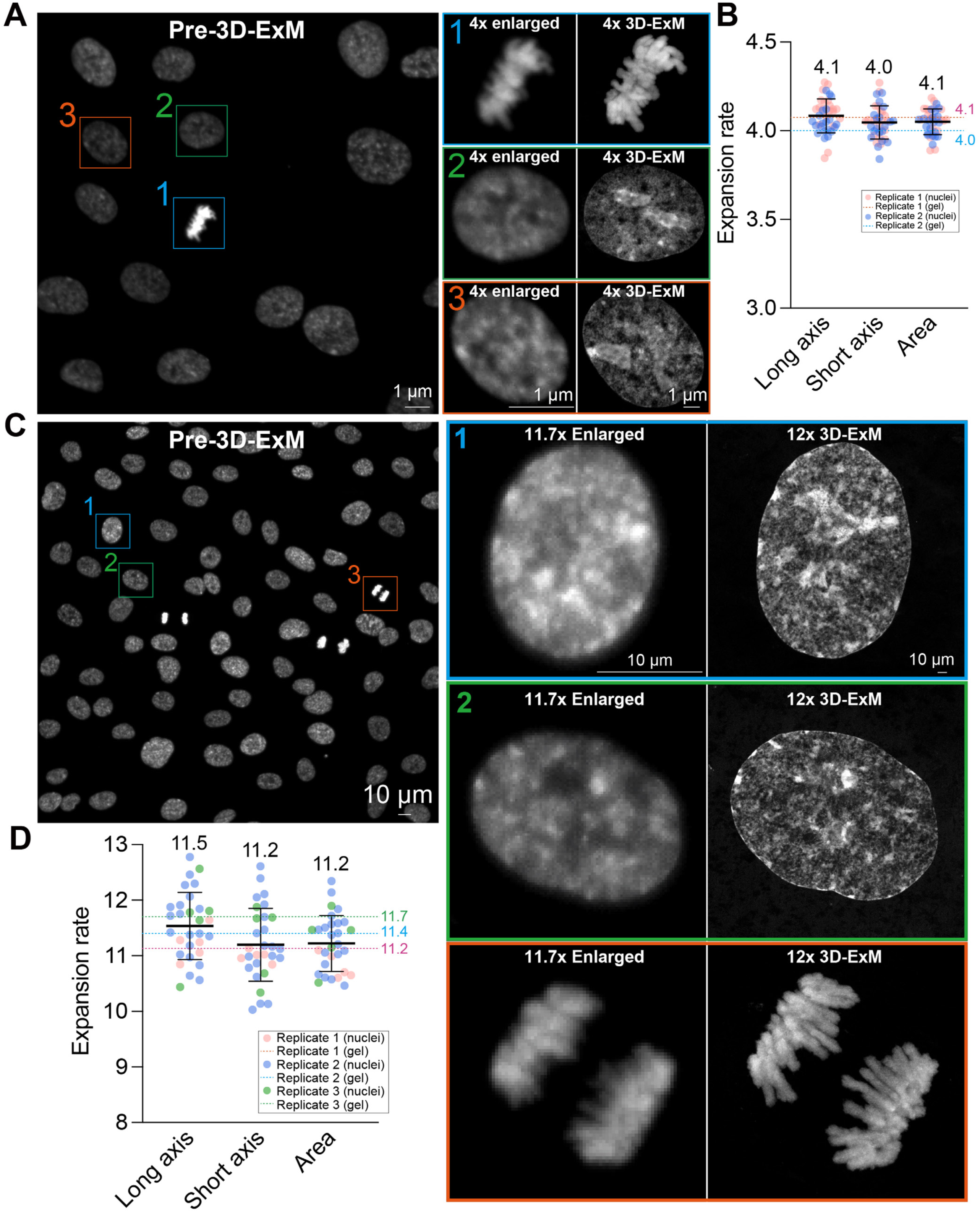

### 12x 3D-ExM enables expansion of 3D organoids model

We investigated whether 3D-ExM could be applied to more complex 3D organoid culture models. Human organoids derived from the MCF-7 breast cancer cell line and primary breast tumor cells were generated, and the expansion fold change of their nuclei was determined following 12x 3D-ExM (**Fig. 4A-B and S3A-B**). The nuclei within the organoids were expanded by approximately 12-fold, as measured by the quantification of their volume and surface area. This significant expansion greatly improved the axial resolution, allowing for the clear identification of individual nuclei within patient-derived organoid (**Fig. S3A**). Collectively, we conclude that 3D-ExM robustly and isotropically expands nuclei in both 2D cell monolayers and 3D organoid culture models.

**Figure 4.**
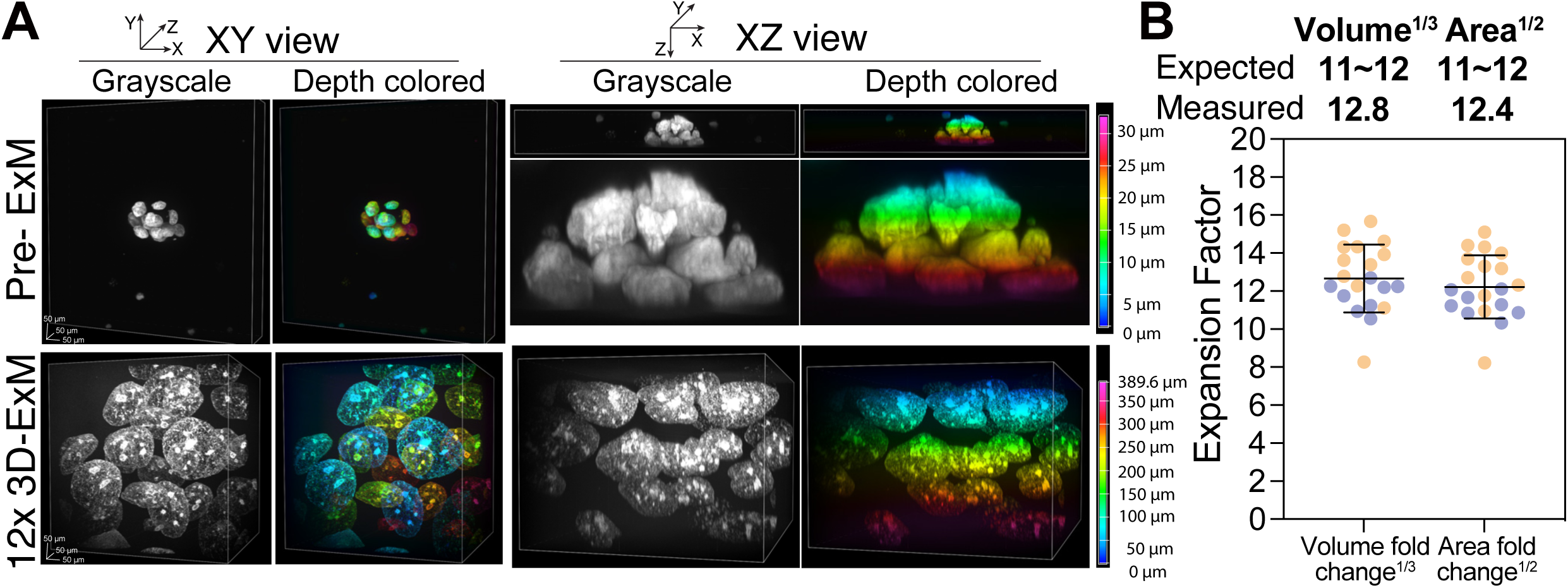

### Validation of achievable resolution of 12x 3D-ExM using cellular rulers

In theory, 4x and 12x 3D-ExM could achieve ∼60 nm and ∼20 nm lateral resolution, respectively, with regular confocal microscopy (based on ∼250 nm lateral resolution of confocal microscopy). To assess the actual performance of 3D-ExM in terms of achievable resolution limits, we first measured the diameter of microtubules (∼25 nm)^24^, a common “cellular ruler”, in interphase PtK2 cells using both confocal and STED microscopy. **Fig. 5A** presents example confocal images of PtK2 cells with fluorescently labeled DNA and microtubules, using the same size of field of view (FOV) before and after expansion (4x and 12x 3D-ExM). We employed line intensity scan to measure the microtubule diameter. As expected, without expansion, the microtubule diameter was ∼250 nm with regular confocal microscopy and ∼130 nm with STED microscopy, respectively (**Fig. 5B-C and S4A**). Using 4x 3D-ExM combined with regular confocal and STED microscopy, the microtubule diameter measured 80 nm and 60 nm, respectively. However, with 12x 3D-ExM, both confocal and STED microscopy yielded a consistent microtubule diameter of ∼28 nm, demonstrating that 12x 3D-ExM combined with regular confocal microscopy can achieve sub-30 nm lateral resolution. Consistent with these findings, 12x 3D-ExM resolved bundled microtubules in both 2D and 3D (**Fig. S4B**). It is noteworthy that the slight difference in the measured microtubule diameter between electron micrographs (25 nm) and 12x 3D-ExM (28 nm) may result from the additional size of the antibody used in our experiments to label alpha-tubulin.

**Figure 5.**
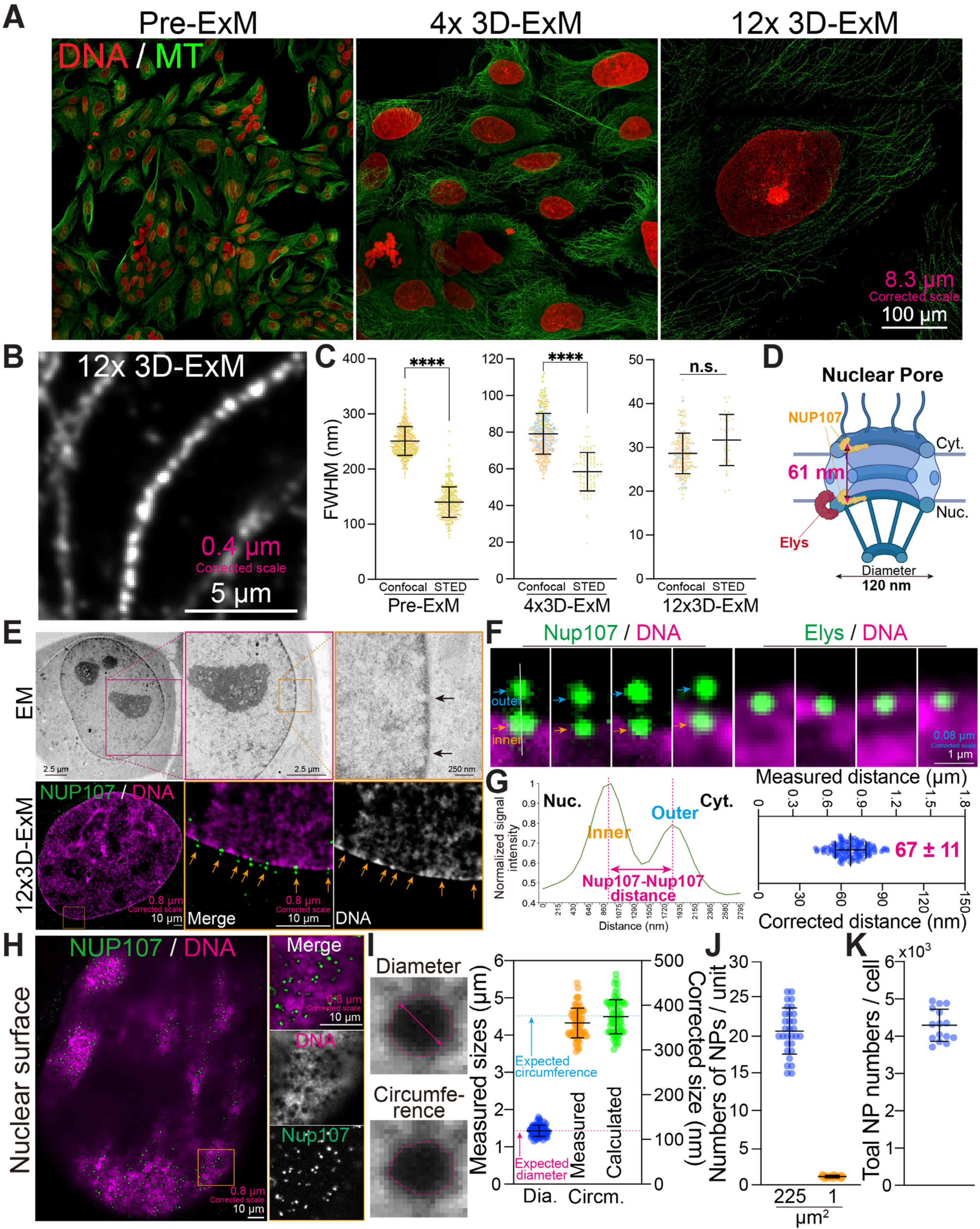

To further validate the resolution achieved by 12x 3D-ExM, we imaged NUP107 and Elys, components of the nuclear pore complex (NPC), also commonly used to evaluate the resolution of imaging techniques^8, 9, 25^ (**Fig. S5A**). NUP107 is a crucial structural component of both the cytoplasmic and nuclear rings of the NPC, with an inter-ring distance of ∼61 nm based on previous cryo-EM data^8^ (**Fig. 5D**). Elys, a nucleoporins, is exclusively located at the nuclear ring of the NPC^9^. Nuclear pores (NPs) were identified as low-electron density regions at the nuclear periphery via transmission electron microscopy (TEM) (**Fig. 5E**). Using 12x 3D-ExM, we observed similar features with relatively low DNA signal levels at the nuclear periphery, overlapping with NP proteins (**Fig. 5E and S5B**). Notably, while Elys appeared as a single dot, NUP107 presented as two foci at each NP, corresponding to the cytoplasmic and nuclear rings of the NPC (**Fig, 5F**). The average corrected distance between NUP107 foci at a single NP was ∼67 nm, consistent with cryo-EM measurements^26^. These results confirm that 12x 3D-ExM readily resolves the two pools of NUP107, ∼60nm apart, within a single NPC, demonstrating a lateral resolution of 12x 3D-ExM with standard confocal microscopy achieves significantly better than the 60 nm (**Fig. 5G)**. Like scanning electron microscopy, 12x 3D-ExM visualizes NPs at the nuclear surface (**Fig. 5H**). Nearly all NPs at the nuclear surface, identified by gaps in the DNA density, colocalized with NUP107 signals (**Fig. 5H and S5C**). The corrected diameter and circumference of NPs were ∼120 nm and ∼380 nm, respectively (**Fig. 5I**), consistent with values from cryo-EM structure^26^. Based on the surface area and NP density, an interphase nucleus of RPE1 cells is estimated to have ∼4300 NPs (**Fig. 5J-K and S5D**), within the range of previous estimates for human nucleui^27, 28^. In summary, 12x 3D-ExM achieves theoretical image resolution with isotropic expansion for both cytoplasmic and nuclear proteins.

### Determination of cellular protein and genomic architectures by 4x and 12x 3D-ExM

We investigated whether 3D-ExM could be employed to discern cellular protein complexes and genomic architectures that are too small to study using conventional fluorescence microscopy. We visualized individual virions of Epstein-Barr virus (EBV), a double-stranded DNA virus and ubiquitous human pathogen of human herpesviruses^29,30^. Previous EM research has determined that the diameter of EBV virions ranged from 100 to 220 nm^31–34^. We utilized the well-characterized iD98/HR1 cell line, an EBV-positive cell line engineered to conditionally enter the lytic stage of EBV’s life cycle, when EBV virions were produced^35, 36^ (See **Methods**). To visualize individual EBV virions, we fluorescently labeled the viral glycoprotein 350 (gp350), the most abundant glycoprotein on the EBV envelope (**Fig. 6A**)^37^. Without 3D-ExM, we observed gp350 signals in lytic cells, but could not resolve individual EBV virions (**Fig. S6**). In contrast, 4x 3D-ExM revealed individual EBV virions, as gp350 foci were observed overlapping with DNA puncta, indicative of encapsidated EBV genomes (**Fig. 6A**). These results showed that 4x 3D-ExM combined with regular confocal microscopy achieved a single EBV virion resolution, thus approaching the 60 nm theoretical resolution limit for 4-fold expansion. Remarkably, 12x 3D-ExM resolves the viral envelope as a ring-like structure surrounding a DNA dot signal, representing the encapsidated EBV genome. In 12x 3D-ExM, the outer and inner diameter of EBV virions were 162 nm and 76nm, respectively, aligning with previous EM data^38^. These findings demonstrate that 12x 3D-ExM has comparable ability to EM, which achieves a resolution sufficient to examine viral architectures (**Fig. 6A and Supplementary Movie 1**).

**Figure 6.**
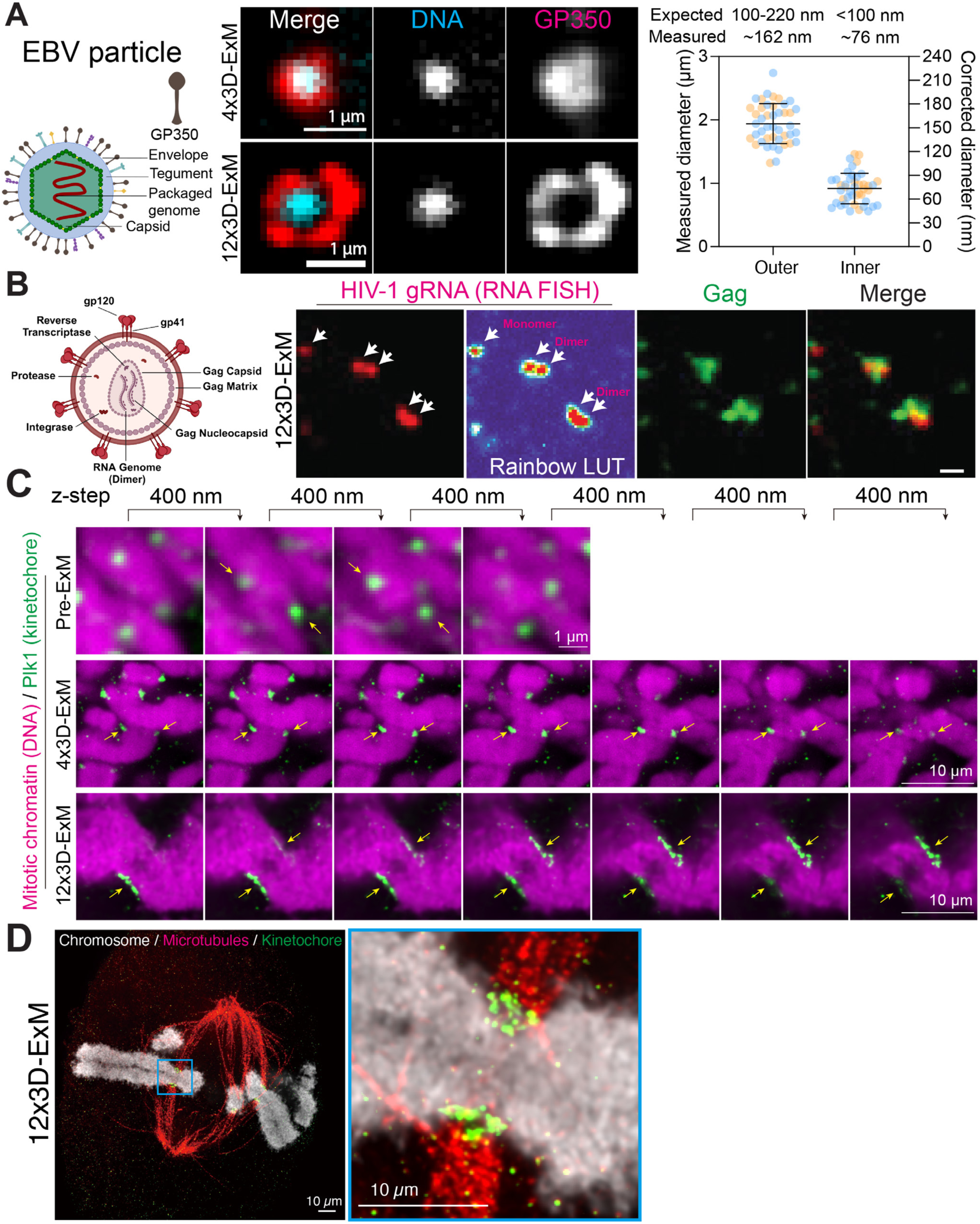

We next asked whether 12x 3D-ExM preserved RNA and enabled the detection of RNA. To this end, we utilized single-molecule fluorescence in situ hybridization (smFISH) to visualize human immunodeficiency virus-1 (HIV-1) unspliced RNA^39^, transcribed from the integration sites within the nucleus containing the wild-type HIV-1 genome (**Fig. S7A**). This custom RNA FISH probe, specific for the gag-pol open reading frame^40^, successfully visualized HIV-1 genome integration site in both pre- and post-12x 3D-ExM (**Fig. S7A-B**). HIV-1 unspliced RNAs are packaged as genome dimers with the viral Gag structural protein during assembly at plasma membrane sites^41, 42^. We performed RNA FISH coupled with Gag immunofluorescence to detect dimerized genomes associated with Gag in virus particles. Our finding revealed that virus particles shed from HIV-1 infected HeLa cells exhibited two fluorescent RNA signal peaks, co-localizing with the Gag signals. This suggests that 12x 3D-ExM can detect both a monomer and a dimer conformation of the HIV-1 unspliced RNA (**Fig. 6B and S7C**). These results demonstrate that 12x 3D-ExM effectively preserves RNA integrity and enables super-resolution imaging of RNA using smFISH. Furthermore, we employed 4x 3D-ExM to visualize the intricate structure of centrioles (**Fig. S8A-C**). Centrioles are a pair of cylindrical structures (mother centriole and procentriole) arranged perpendicular to each other^43^. Unlike the procentriole, the mother centriole is distinguished by the presence of distal appendage at its distal end^43^. To examine these structures, we labeled CEP164 (a marker for distal appendage) and acetylated-tubulin (a marker for the centriole wall) in cold-treated RPE1 cells. The 4x 3D- ExM revealed the cartwheel structure of the mother centriole, showcasing a universal 9-fold radial symmetry from the top view perspective. Additionally, the distal appendage was observed as a protrusion at one end of the mother centriole from the side view perspective (**Fig. S8A**). The measured diameter of distal appendage ring and the length of centrioles obtained through 4x 3D-ExM were consistent with previous measurements using other super-resolution imaging methods^44–46^ (**Fig. S8B-C**).

The kinetochore is a macro-molecular protein complex that assembles on centromeres, serving as a microtubule attachment site and orchestrating chromosome movements during mitosis^47^. The human kinetochore displays a plate-like structure, approximately 300 nm length and depth with 50 nm thickness, as observed by EM^48, 49^. Due to the lateral PSF and 3D orientation of kinetochores, they usually appear as puncta in conventional fluorescence microscopy rather than rectangle plates (**Fig. 6C, top,** Plk1 antibody as a kinetochore marker^50^). While the lateral PSF does not interfere when the objects are larger than 250 nm in length, the axial PSF significantly overestimates the size, even if the axial length exceeds 1 µm^2^. Supporting this, kinetochore signals prior to 3D-ExM appear in 4 separate optical sections spaced at 400 nm depth, overestimating the size to more than 1.2 µm depth compared to the ∼300 nm size determined by EM (**Fig. 6C, top and S9A**). With 4x 3D-ExM, kinetochores appeared more rectangular, though still not matching the plate-like structures seen in EM^48, 49^.. The depth was also significantly overestimated, showing ∼3.2 µm optically (∼800 nm corrected) in 4x 3D-ExM (**Fig. 6C, middle and S9B**). These discrepancies are likely due to the combined effects of axial PSF and the 3D orientation of kinetochores^2^. In contrast, kinetochores in 12x 3D-ExM exhibited clear plate-like structures with an expected depth of ∼3 µm optically (∼250 nm corrected) (**Fig. 6C, bottom and S9C**). The 12x 3D-ExM technique enables visualization of the kinetochore-microtubule interface with a resolution previously unattainable through light microscopy **(Fig. 6D)**. Collectively, our data illustrate that 12x 3D-ExM provides sufficient resolution to minimize the effects of axial PSF and object orientation in 3D on the shape and size of protein architectures below the diffraction limit, such as kinetochores, allowing for accurate 2D/3D measurements of these cellular structures.

### Practical applications of 12x 3D-ExM

12x 3D-ExM technique achieves unprecedented resolution through a single expansion process. We illustrated its capabilities with two practical examples, serving as proof of principle. Aneuploidy, characterized by the gain or loss of chromosomes due to mitotic errors, is a crucial aspect of cancer and influences therapeutic outcomes^51^. To accurately determine the karyotype of cells and evaluate the pattern and extent of aneuploidy, the chromosome spread technique is commonly employed. This method involves swelling and bursting of cells in hypotonic solution. However, this process frequently results in chromosome overlap, which can lead to significant errors in quantification and identification^52^. Recent studies have attempted to map chromosomes in intact cells using serial block-face scanning electron microscopy (SEM)^53, 54^. However, these efforts have only successfully identified a subset of chromosomes in nocodazole-treated cells and were unable to perform quantifications in multiple cells due to the time-consuming nature of the method. 12x 3D-ExM is expected to achieve resolution comparable to or better than serial block-SEM^54^, with the added advantage of specifically labeling target molecules. To ascertain if the complete karyotype of PtK2 cells could be accurately identified without using chromosome spreads, we utilized immunofluorescence to label kinetochores and DNA in asynchronous PtK2 cells, which possess 14 chromosomes^55^. As anticipated, this method allowed us to distinctly visualize each chromosome in intact metaphase cells (**Fig. 7A and Supplementary Movie 2**). Furthermore, we successfully identified all chromosomes in these cells by analyzing chromosome size and relative kinetochore location along the chromosome (**Fig. 7A and S10A-C**). Using the same approach, we determined the aneuploidy status in an intact single cell (**Fig. 7A and Supplementary Movies 3-4**). We identified aneuploid cells by simply counting the number of chromosomes in an intact mitotic cell, pinpointing which chromosomes were lost or gained without the need for chromosome spreading, FISH, or the averaging of multiple cell quantifications. These results demonstrate that 12x 3D-ExM can accurately determine the complete karyotype of an intact mitotic cell on a single-cell basis. Additionally, this method enables the exploration of the entire chromosomal geometric information at any stage of mitosis, which is likely crucial for understanding the mechanisms underlying chromosomal instability (CIN)^56^.

**Figure 7.**
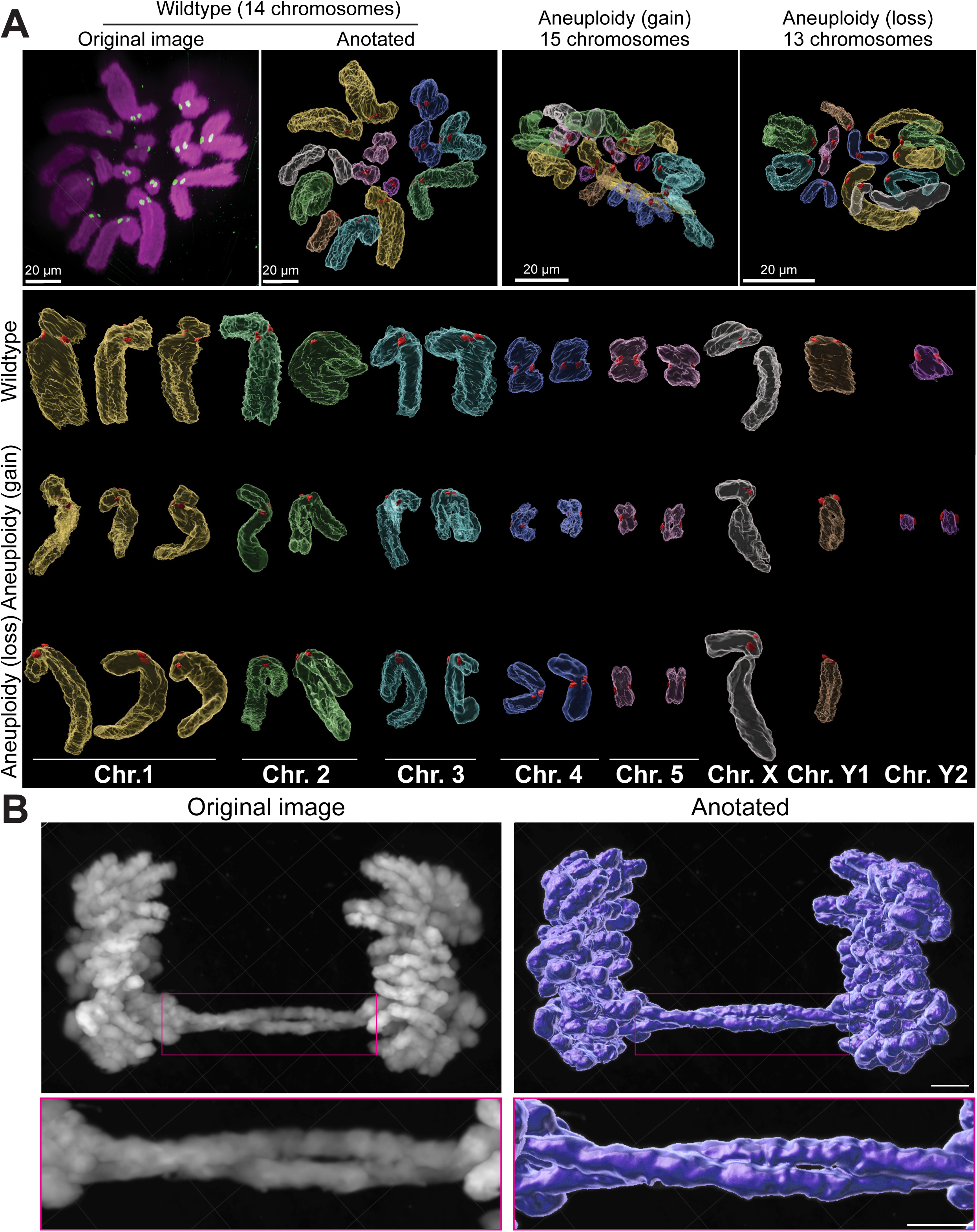

Chromosome bridge, a type of mitotic error, is defined by chromatin linkages between sister chromatids during anaphase. These bridges often result in the formation of micronuclei, which lead to chromothripsis and chromosomal instability (CIN)^57^. They often arise from DNA damage and cohesion defects^58^. An example 12x 3D-ExM image of HeLa cells with a chromosome bridge was shown in **Fig. 8B**. Unexpectedly, this 12x 3D-ExM image revealed that the chromosome bridge could be comprised of two intertwined sister chromatids (**Fig. 8B**). This finding underscores the capability of 12x 3D- ExM to uncover previously invisible chromosomal architectures, potentially leading to novel discoveries.

## Discussion

Fluorescence microscopy was invented over 100 years ago. Since then, both fluorescence microscopes and fluorophores, such as dyes and proteins, have been continuously improved, expanding their applications across many research fields. To address the need for studying subcellular structures below the resolution limit of conventional light microscopy, various super-resolution imaging technologies have been developed. However, these methods often require expensive equipment, special dyes or reagents, and complex post-image processing, limiting their accessibility. In this study, we demonstrate 3D-ExM, a specimen-based super-resolution microscopy approach modified from traditional ExM, which achieves robust < 30 nm lateral and < 50 nm axial resolution. This method surpasses common optical- or post-image processing-based super-resolution techniques and does not require specialized equipment, oxygen removal, or iterative expansion processes. Additionally, 3D-ExM can be combined with common super-resolution microscopes for further improved resolution. Featuring an affordable and user-friendly protocol, 3D-ExM makes super-resolution microscopy accessible to all researchers, facilitating nanoscale discoveries in various research fields.

The primary challenges in ExM technologies include the complex protocols required for achieving expansions greater than 4-fold, maintaining isotropy in 2D/3D expansions across all organelles, and preserving cellular structures. In this study, we demonstrated that the interphase nucleus expands uniformly according to gel expansion rates, using correlative pre- and post-3D-ExM methods. Additionally, population measurements confirmed that 3D-ExM consistently achieves 3D isotropic expansion of the nucleus in both 2D and 3D culture systems. Furthermore, 3D-ExM effectively preserves protein architectures, chromatin, and RNA, as evidenced by the visualization of EBV virions, HIV-1 genomic RNA, kinetochores, mitotic chromosomes, centrosomes, microtubules, and nuclear pores. The 4x and 12x 3D-ExM methods are among the most comprehensively validated techniques and are applicable to a broad range of research fields, including nuclear and chromosome research.

ExM techniques are continuously advancing to improve resolution. Achieving single-protein resolution with standard confocal microscopy would require an expansion greater than 12-fold, ideally between 50-100-fold. Inspired by iterative expansion microscopy (iExM)^17^ and its recent applications to chromatin (chromExM)^59^, combining 4x and 12x 3D-ExM or repeating 12x 3D-ExM may achieve approximately 50- and 150- fold expansion, respectively. Collectively, 3D-ExM is an affordable super-resolution imaging tool accessible to researchers without specialized equipment, enabling nanoscale visualization and quantifications of protein and genomic architectures.

## Methods

### Reagents

#### Immunostaining

- Phosphate buffered saline (PBS): Sigma-Aldrich, cat. no. P3813
- Paraformaldehyde (PFA): Sigma-Aldrich, cat. no. P6148
- Nonidet P-40 Substitute (NP40): SCBT, cat. no. sc-29102
- Bovine Serum Albumin (BSA): Sigma-Aldrich, cat. no. A2153

#### Crosslinking

- 70% Glutaraldehyde solution (GA): Sigma-Aldrich, cat. no. G7776

#### 4x 3D-ExM gel

- Phosphate buffered saline (PBS): Sigma-Aldrich, cat. no. P3813
- Sodium Chloride (NaCl): Fisher BioReagents, cat. no. BP358-212
- Acrylamide: Sigma-Aldrich, cat. no. A9099
- N,N’-Methylenebisacrylamide (MBAA): Sigma-Aldrich, cat. no. M7279
- Sodium acrylate (SA): Sigma-Aldrich, cat. no. 408220
- 4x Monomer solution (4x 3D-ExM MS): 1x PBS, 2 M NaCl, 2.5% w/v Acrylamide, 0.15% w/v MBAA, 8.6% w/v SA
- Ammonium persulfate (APS): Sigma-Aldrich, cat. no. A3678
- N,N,N’,N’-Tetramethylethylenediamine (TEMED): Sigma-Aldrich, cat. no. T7024
- 4x Gelling solution (4x 3D-ExM GS) = 95% 4x MS + 1 % of ddW + 2% of 10% APS + 2 % of 10% TEMED

#### 12x 3D-ExM gel

- N,N-Dimethylacrylamide (DMAA): Sigma-Aldrich, cat. no. 274135
- Sodium acrylate (SA): Sigma-Aldrich, cat. no. 408220
- 12x Monomer solution (12x 3D-ExM MS): 1.335 g DMAA + 0.32 g SA + 2.85 ml of ddH_2_O
- Potassium persulfate (KPS): Sigma-Aldrich, cat. no. 379824
- N,N,N’,N’-Tetramethylethylenediamine (TEMED): Sigma-Aldrich, cat. no. T7024
- 12x Gelling solution (12x 3D-ExM GS): 90% of MS + 10% of 0.036 g/ml KPS + 0.8 μl of TEMED

#### Protein digestion

- Triton X-100: Sigma-Aldrich, cat. no. T9284
- Sodium dodecyl sulfate (SDS): Roche, cat. no. 1667289
- TTSDS buffer: 1x TAE, 0.5% Triton X-100, 1% SDS, ddH_2_O
- Proteinase K (ProK): Thermo Scientific, cat. no. EO0492
- Digestion solution (DS): 16 U/ml ProK / TTSDS buffer

### Cell Culture

Human HeLa, PRE1, T47D, MCF7, and rat kangaroo PtK2 cells were originally obtained from the American Type Culture Collection (ATCC, Manassas, VA, USA). HeLa (DMEM High glucose, Cytiva Hyclone; SH 30243.01), RPE (DMEM F12, Cytiva Hyclone; SH 3026101) and T47D (RPMI, Fisher; SH 30255.01) cells were grown as monolayer cultures on 12-mm # 1.5 circular coverslips in their corresponding growth media supplemented with 1% penicillin-streptomycin, 1% L-glutamine, and 10 % fetal bovine serum under 5% CO_2_ at 37°C in an incubator. Ptk2 cells were cultured in EMEM media (Gibco, 12492013) supplemented with 20% FBS and 1% penicillin-streptomycin, 1% L-glutamine, under 5% CO_2_ at 37°C. MCF7 (gift from Andreas Friedl) or patient derived cells were cultured as 3D spheroids in a 1:1 mixture of DMEM/High Glucose media and Matrigel (Corning; 354230). Cells were passaged as 40 µL droplets/well in a 24-well plate, nourished with 500 µL media. For imaging, spheroids were dissociated to small clusters with trypsin, pelleted and re-suspended in a 1:1 mixture of culture media and Matrigel. Patient tissue was collected with informed consent from all patients in accordance with Health Insurance Portability and Accountability Act (HIPAA) regulations, and all studies were approved by the IRB at the University of Wisconsin–Madison (IRB# UW14035, approval no. 2014-1053). Eligible patients were planned for ultrasound biopsy meeting certain criteria determined by the Diagnostic Radiologist. All subjects provided written informed consent. For imaging, spheroids were dissociated to small clusters with trypsin, pelleted and re-suspended in a 1:1 mixture of culture media and Matrigel. They were plated as 10 µL droplets onto 12-mm # 1.5 circular coverslips in a 24-well plate, nourished with 500 µL media until fixation.

### Antibodies and Dyes

Following primary antibodies were used in this study: mouse anti-Plk1 antibody (Santa Cruz Biotech, sc17783, 1:100), mouse anti-alpha Tubulin antibody (Sigma, DM1a, T6199, 1:200), mouse anti-Nup107 antibody (Abcam, ab24609, 1:500), rabbit anti-Elys (SinoBiological, 205696-T10, 1:100), anti-gp350 antibody (SinoBiological, 40373, 1:100), mouse monoclonal Gag antibody (183-H12-5C; 1:1,000 dilution) from Bruce Chesebro and obtained from the NIH AIDS Research and Reference Reagent Program (Bethesda, MD, USA)^60^. The secondary antibodies used are Rat anti-mouse IgG2a-biotin (Fisher, 13-4210-80, 1:100), Rat anti-mouse IgG1-biotin (Fisher, 13-4015-80, 1;100), and Minimal cross-react Alexa 488 conjugated antibodies against rabbit, mouse, and goat IgG (JacksonImmuno, 711-545-152, 111-545-144, 715-545-150, 115-545-146, 805-545-180, 1:300). To label biotinylated secondary antibodies, fluorescently labeled streptavidin (Alexa fluor 488 (S32354), 546 (S11225), or 594 (S32356), Thermofisher (1:200)) was used. For DNA staining, DAPI ((4’,6-diamidino-2-phenylindole, Thermo, D1306) or Draq5 (Thermo, 62252) were used.

### Fixation and staining for 3D-ExM (pre-embedding staining)

Cells are fixed with 3% PFA in PHEM (60 mM PIPES, 27.2 mM HEPES,10 mM EGTA, 8.2 mM MgSO_4_) for 15 min at 37°C followed by 3 times of PBS wash. Cells used for pore-C experiments were fixed with 1% PFA in PHEM. For microtubule staining, cells are fixed with Glutaraldehyde (GA) solution (0.8% GA, 1% Triton X-100, 3% PFA, PHEM buffer) and quenched by NaBH_4_. Fixed cells are permeabilized with 0.5% NP40 at room temperature (RT) for 15 min. Cells are incubated with BSA at RT for 30 min. Then, cells are incubated with primary antibody solution (antibodies are listed in Antibodies) and secondary antibody solution at 37°C in a humidified chamber. If biotin conjugated secondary antibodies are used, cells are incubated with fluorophore-conjugated streptavidin at 37°C in a humidified chamber. Crosslink is performed with 2% GA / PBS at RT for 12-16 hr (overnight). Prepare a glass slide with a square mold on top. Transfer the stained coverslip into the mold with the cell side facing up.

### 4x 3D-ExM (Continue from Fixation and staining)

Add 4x 3D-ExM monomer solution (MS) to infiltrate the cells at 4°C for 30 min. Replace 4x 3D-ExM MS with 4x 3D-ExM gelling solution (GS) and incubate the cells at 37°C in a humidified chamber for 1 hr to form the gel. Add digestion solution (DS) onto the gel with a piece of parafilm on top and incubate the gel at 37°C for 3 hr in a humidified chamber. Transfer the gel from the coverslip to a large dish filled with ddH_2_O. Incubate the gel in ddH_2_O for 3 hr and replace ddH_2_O every 30 min to allow for gel expansion.

### 12x 3D-ExM (Continue from Fixation and staining)

Add 12x 3D-ExM MS to infiltrate the cells at RT for 10 min. Replace 12x 3D-ExM MS with 12x 3D-ExM GS and incubate the cells at RT in a humidified chamber for 2 hr to form the gel. Place a piece of parafilm on top of the mold to prevent air exposure. Add DS onto the gel with a piece of parafilm on top and incubate the gel at 37°C for 24 hr (overnight) in a humidified chamber. Transfer the gel from the coverslip to a large dish filled with ddH_2_O. Incubate the gel in ddH_2_O for at least 20 hr (overnight) and replace ddH_2_O every 1 hr during the first 6 hours to allow for gel expansion.

### Generation and titration of HIV-1 Vif-/Vpr-CFP virus

Human embryonic kidney (HEK) 293T cells at 30-40% confluency were transfected with 10 ug plasmid DNA (9ug plasmids encoding a biosafe, single-round HIV-1 Env-/Vif-/Vpr-/Nef-/CFP reporter virus with 1ug plasmid encoding the G protein from vesicular stomatitis virus (VSV-G) for virion pseudotyping) in 10cm dishes using polyethylenimine (PEI; catalog no. 23966; Polysciences Inc., Warrington, PA, USA). Culture media were replaced at 24 hours post-transfection. At 48 hours post-transfection, virion-containing supernatant was harvested and filtered.

### HIV-1 infection, fluorescence in situ hybridization (FISH), and immunofluorescence

RNA-FISH method for HIV-1 genome is described in previous study^39^. HeLa cells were plated on 12mm circular coverslips (#1.5 thickness) in 12-well plates and allowed to grow to 30-40% confluency prior to infection. Cells were infected with 500uL of the above HIV-1 Env-/Vif-/Vpr-/Nef-/CFP reporter virus using DEAE-Dextran (catalog no. 9064-91-9; Millipore Sigma, Darmstadt, Germany) at a concentration of 6 mg/mL. At 24 hours post-infection, culture media was replaced, and at 48 hours post-infection cells were washed with PBS and fixed in 3.7% formaldehyde in PBS. Cells were permeabilized with 70% ethanol for at least 1 hour at 4°C. Custom Integrated DNA Technologies (IDT) probes were designed against NL4-3 HIV-1 unspliced (US) RNA specific for the gag-pol open reading frame (nucleotides 386 to 4614) and containing a biotin modification of the 5’-ends of each probe (48 probes total). Cells were hybridized with the Gag/Gag-Pol IDT Biotin RNA FISH probe set using Stellaris FISH (Biosearch Technologies, Inc.) buffers and following the Stellaris FISH instructions available online at www.biosearchtech.com/stellarisprotocols. Immunofluorescence was carried out after hybridization of the biotin probes. Cells were washed with PHEM buffer and blocked in 0.1% BSA/PHEM solution for 30 minutes at 37°C. Primary antibody to Gag p24^60^ (1:100) and streptavidin secondary antibody (1:100) were diluted in blocking buffer and incubated for 3 hours at 37°C. Cells were washed in PHEM prior to incubation in secondary Gag antibody (Goat anti-Mouse IgG (H+L) Cross-Adsorbed Secondary Antibody, Alexa Fluor™ 594 from Thermo Fisher Scientific, catalog # A-11005, RRID AB_2534073; 1:100) diluted in blocking buffer for 2 hours at 37°C. An additional Gag secondary/tertiary antibody (Donkey anti-Goat IgG (H+L) Cross-Adsorbed Secondary Antibody, Alexa Fluor™ 594 from Thermo Fisher Scientific, catalog # A-11058, RRID AB_2534105; 1:100) was diluted in blocking buffer and incubated with cells for 1 hour at 37°C. Finally, cells were washed three times in PHEM buffer prior to 12x 3D-ExM procedure.

### Imaging

Nikon Ti2 stand equipped with Yokogawa SoRa CSU-W1 spinning disc confocal, a Yokogawa uniformizer, Hamamatsu Orca Flash4 cameras, and a high-power laser unit (100 mW for 405, 488, 561, 640 nm wavelength). Z-stack images were acquired at a step of 0.1∼0.4 μm (mostly 0.4 µm for 3D-ExM images) by Nikon NIS element software (version 5.20). Plan Apo VC 60x water objective (NA 1.2), Plan Apo 40x or 25x Silicon objectives, and Plan Apo 100x oil (NA 1.5) were used. A house-made gel chamber is used for 3D imaging.

### Statistics

The data is represented as the mean ± standard deviation (s.d.). Welch’s t-test was used to compare the means between two populations. p < 0.05 was considered statistically significant. All quantification were performed at least two biological replicates (most of data were three replicates). Sample numbers and numbers of replicates were stated in each figure legend.

## Acknowledgement

We would like to thank Drs. Nathan Claxton, Hiroshi Nishida, Yoshitaka Sekizawa, the University of Wisconsin Optical Imaging Core, Yokogawa Electrical Corporation, Nikon Japan, and Nikon USA for critical equipment and technical support. We also would like to thank Drs. William Sugden, Beth Weaver, Robert Lera, Emily Kaufman, Yu-Lin Chen and Rebeca Garcia-Varela for the critical suggestions and experimental support. Part of this work is supported by Wisconsin Partnership Program, the University of Wisconsin-Madison Office of the Vice Chancellor for Research with funding from the Wisconsin Alumni Research Foundation, start-up funding from University of Wisconsin-Madison SMPH, UW Carbone Cancer Center, and McArdle Laboratory for Cancer Research, and NIH grant R35GM147525 (to A.S.), R01GM131068 and R01CA234904 (to M.E.B), U54AI170660, R01AI110221, and P01CA022443 (to N.S.), and P20GM104360, UND COBRE pilot genomics awards, and startup fund and Dean’s fund from UND (M.T.), and JSPS fellowship (M.S.).

## Author contribution

E.R. and R.N. initiated the project. E.R., under the guidance of A.S., developed the method for 12x gel polymerization. R.N. and Y.C., with assistance from J.L. and E.R., conducted the majority of the experiments. Q.R. and J.L. were responsible for Fig. 6A and S6, while SL.L. and N.S., with assistant from R.N., conducted Fig. 6B and S7. Y.C. performed all the work for Fig. 1B, 3, 5A, 7A, S4, S8, and S10. A.S. conceptualized and supervised the entire project, contributing pivotal ideas and designing the experiments. ME.B., N.S., and M.T. offered valuable suggestions and oversaw the experiments conducted by R.N. and SL.L. Y.C. and A.S. prepared the initial manuscript draft, with contributions from M.T., ME.B., and R.N. All authors reviewed and contributed to the manuscript’s refinement.

## Competing Financial Interests

A. Suzuki, ME. Burkard, R. Norman, and E. Recchia declare partial ownership (5% each) of US Provisional Patent application US-2022-0074829 titled “Optimized Economical and Modulatable Isotropic Expansion Microscopy”.

The authors declare no further conflict of interests.

## Notes

### Competing Interest Statement

The authors have declared no competing interest.

